# Principal component based adaptive association test of multiple traits using GWAS summary statistics

**DOI:** 10.1101/269597

**Authors:** Bin Guo, Baolin Wu

## Abstract

Genetics hold great promise to precision medicine by tailoring treatment to the individual patient based on their genetic profiles. Toward this goal, many large-scale genome-wide association studies (GWAS) have been performed in the last decade to identify genetic variants associated with various traits and diseases. They have successfully identified tens of thousands of disease-related variants. However they have explained only a small proportion of the overall trait heritability for most traits and are of very limited clinical use. This is partly owing to the small effect sizes of most genetic variants, and the common practice of “testing association between one trait and one genetic variant at a time” in most GWAS, even when multiple related traits are often measured for each individual. Increasing evidence suggests that many genetic variants can influence multiple traits simultaneously, and we can gain more power by testing association of multiple traits simultaneously. It is appealing to develop novel multi-trait association test methods that need only GWAS summary data, since it is generally very hard to access the individual-level GWAS phenotype and genotype data.

Most existing GWAS summary data based association test methods have relied on ad hoc approach or crude Monte Carlo approximation. In this paper we develop rigorous statistical methods for efficient and powerful multi-trait association test. We develop robust and efficient methods to accurately estimate the marginal trait correlation matrix using only GWAS summary data. We construct the principal component (PC) based association test from the summary statistics. PC based test has optimal power when the underlying multi-trait signal can be captured by the first PC, and otherwise it will have suboptimal performance. We develop an adaptive test by optimally weighting the PC based test and the omnibus chi-square test to achieve robust performance under various scenarios. We develop efficient numerical algorithms to compute the analytical p-values for all the proposed tests without the need of Monte Carlo sampling. We illustrate the utility of proposed methods through application to the GWAS meta-analysis summary data for multiple lipids and glycemic traits. We identify multiple novel loci that were missed by individual trait based association test.

All the proposed methods are implemented in an R package available at http://www.github.com/baolinwu/MTAR. The developed R programs are extremely efficient: it takes less than two minutes to compute the list of genome-wide significant SNPs for all proposed multi-trait tests for the lipids GWAS summary data with 2.5 million SNPs on a single Linux desktop.

## 1 Introduction

Genetics hold great promise to precision medicine that tailors the treatment to the individual patient by considering their genetic profiles. Many large-scale genome-wide association studies (GWAS) performed in the past decade have successfully identified thousands of common genetic variants associated with various traits and diseases (Visscher *et al.*, 2017). But in total they have explained a small proportion of the overall heritability (Manolio *et al.*, 2009) and are of limited clinical use. It is expected that many more genetic variants remain to be identified. These GWAS are primarily based on the paradigm of “single trait single variant association test” even with multiple traits measured for each individual. There is increasing evidence showing that genetic variants can influence multiple traits simultaneously, and we can gain great power by studying multiple traits simultaneously (see, e.g., Ferreira and Purcell, 2009; Tang and Ferreira, 2012; O’Reilly *et al.*, 2012; Stephens, 2013; Maity *et al.*, 2012; Seoane *et al.*, 2014; Galesloot *et al.*, 2014; Zhu *et al.*, 2015; Wu and Pankow, 2016; Broadaway *et al.*, 2016). Ideally we can reanalyze those existing GWAS data published in the last decade using the multi-trait association test approach to identify more novel genetic variants. However due to privacy concerns and various logistical considerations, it is generally very hard to access the individual-level GWAS phenotype and genotype data, which creates a barrier to further mine these existing data to extract more information.

Nevertheless most published GWAS have made the association test summary statistics publicly available. They include, e.g., the minor allele frequency (MAF), the estimated effect sizes with their standard errors, and significance p-values for each single nucleotide polymorphism (SNP) analyzed in a GWAS. One viable solution is to develop new association test methods that depend on only these GWAS summary data (Pasaniuc and Price, 2017). For example, for the single variant based association test, the GWAS meta-analysis (Evangelou and Ioannidis, 2013) is typically conducted based on the summary statistics, which can be as efficient as analyzing individual-level data across all studies (Lin and Zeng, 2010). Similar methods have been developed for meta-analysis of the rare variant set association across studies (Hu *et al.*, 2013; Lee *et al.*, 2014). For joint association test of a single variant with multiple traits, Stephens (2013) and Zhu et *al.* (2015) proposed methods using only individual GWAS summary statistics and GWAS meta-analysis summary results. The key insight of these approaches is that for a single variant, the summary Z-statistics across different traits share the same correlation as the traits (Stephens, 2013). This has motivated the widely used estimate based on the sample correlations of genome-wide summary Z-statistics, which implies that ideally only independent SNPs should be used in calculating the sample correlation matrix, and further nearly all of them are null SNPs. Therefore some forms of variant filtering based on the linkage disequilibrium (LD) and P-value are needed. This approach can produce significantly biased estimates of trait correlations, and lead to increased false positives in the downstream association test. In this work, we conduct thorough investigation of the impact of trait correlation estimation, and develop a much more accurate and efficient estimate than the empirical correlation matrix. We further implement the proposed methods in a publicly available R package.

The rest of the paper is organized as following. We introduce the proposed adaptive multi-trait association test in Section 2. Section 3 is devoted to simulation studies and comprehensive analyses of GWAS meta-analysis summary data for multiple lipids and glycemic traits. We end the paper with a discussion in Section 4. All technical derivations are delegated to the Supplementary Materials.

In this work, we make several contributions to studies of multi-trait association test using GWAS summary data. First, we propose a robust LD score regression to estimate the trait correlation, which performs much better than the existing approach. Second, although not a major focus of current paper, the proposed robust LD score regression also provides a good approach to estimating genetic correlation using GWAS summary data. Third, we develop powerful, efficient and robust adaptive multi-trait association test methods based on the summary statistics. The proposed methods are extremely scalable to genome-wide association test: we can quickly and accurately compute analytical p-values without the need of Monte Carlo approximation. We have implemented all our proposed methods in an R package.

## 2 Methods

Throughout the following discussion, we mainly focus on analyzing summary association statistics for multiple traits from a common cohort (either a single GWAS or a common GWAS meta-analysis). The proposed methods can be readily extended to partially overlapped GWAS (see supplementary materials section 1.3 for technical details). We take a two-step approach to conduct the summary statistics based multi-trait association test: first, we estimate the trait correlation matrix using the summary statistics of all SNPs; second, with the estimated correlation matrix, we test the association of each SNP with multiple traits using developed methods detailed as follows.

### 2.1 Estimate trait correlation from summary statistics

When testing the marginal association of a SNP with multiple continuous traits, its summary Z-statistics across traits asymptotically follow the multivariate normal distribution with correlation, denoted as ∑ =(*ρ*_*ik*_), equal to the trait correlation matrix (see Stephens, 2013, Zhu *et al.*, 2015, and supplementary materials section 1.1). This has motivated the commonly used approach of empirically estimating ∑ using the sample correlation of genome-wide summary Z-statistics, which implies that ideally only independent SNPs should be used in calculating the sample correlation matrix (therefore some LD pruning is required), and further nearly all of them are null SNPs (hence some P-value filtering is required). For traits of polygenic nature, as shown in Bulik-Sullivan et *al.* (2015a), in addition to the variant LD and trait dependence, both the trait heritability and genetic correlation contribute to the correlation of summary Z-statistics.

Consider a cohort of *N* samples with *K* quantitative outcomes, and assume we have the summary Z-statistics (*z*_1*j*_, …, *z*_*Kj*_) for testing the marginal association of each trait with SNP *j* = 1,…, *M*. We can check that

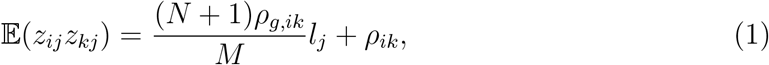

where *ρ*_*ik*_ is the correlation between the *i*-th and *k*-th outcomes, *l*_*j*_ denotes the LD score for SNP *j* (sum of its LD *r*^2^ with all other SNPs), *ρ*_*g*,*ik*_ measures the between trait genetic covariance. For traits of polygenic nature, there could be significant genetic heritability and covariance, leading to potentially non-ignorable *ρ*_*g*,*ik*_. Naively using the sample correlation of summary Z-statistics will then lead to biased estimates of ∑. We can obtain more accurate estimates of ∑ by regressing the pairwise product of summary statistics (*z*_*ij*_*z*_*kj*_) on *l*_*j*_.

To reduce the impact of large summary statistics, it has been a common practice to filter out those large summary statistics (Bulik-Sullivan *et al.*, 2015a), which however is less efficient and can potentially lead to biased estimation. We propose to use a robust linear regression to incorporate all summary statistics: instead of removing those large summary statistics, we minimize their absolute differences in the regression.

### 2.2 Multi-trait association test

For simplicity of notation, denote the estimated correlation matrix as ∑, and consider a single SNP with the across-trait summary statistics vector *Z* = (*z*_1_, …, *z*_*K*_)^*T*^. We can construct the following principal component (PC) based association test statistic, *B* = (*Z*^*T*^*u*_1_)^2^/*d*_1_, where *d*_1_ is the largest eigenvalue and *u*_1_ is the corresponding eigenvector computed from ∑. We can check that *B* asymptotically follows the 1-DF chi-square distribution. *B* performs well when the first PC captures majority of the association signals across multiple traits. Alternatively we can use the omnibus test, *Q* = *Z*^*T*^∑^−1^*Z*, which can detect any deviation from the null. *Q* asymptotically follows the K-DF chi-square distribution. *Q* generally has robust performance under different disease models. To obtain optimal test power, we can consider the adaptive test (denoted as AT) based on their weighted average, *Q*_*ρ*_ = (1 − *ρ*)*Q* + *ρB*, and use the minimum p-value, *T* = min_*ρ*_ *P*(*Q*_*ρ*_), as the test statistic. Here *P*(*Q*_*ρ*_) denotes the p-value of *Q*_*ρ*_. We can quickly and *exactly* compute the analytical p-value of AT (see supplementary materials section 2 for technical details).

## 3 Simulation study

### 3.1 Marginal trait correlation estimation

We first evaluate the estimation of between trait correlations using the GWAS summary statistics. We consider three continuous polygenic outcomes and set the genetic and environment correlation matrices based on the lipids data as ∑_*g*_ = (0.08, −0.59, 0.39), ∑_*e*_ = (−0.19, 0.77, −0.50) (here we only list the three pairwise correlations; see supplementary materials section 3.1 for details). We assume the same heritability *h*^2^ for three traits and the marginal trait correlation matrix ∑ is then *h*^2^∑_*g*_ + (1 − *h*^2^)∑_*e*_.

To mimic a true genome-wide LD structure in the simulation, we use the 9713 European GWAS samples from the Atherosclerosis Risk in Communities Study (ARIC; dbGaP: phs000280.v3.p1) and consider approximately 1.2 million HapMap3 common SNPs. We select *M* = 6100 independent causal SNPs that are approximately 400KB apart. We further divide these causal SNPs into two sets: *M*_*b*_ of them have genetic covariance *h*^2^∑_*g*_/(2*M*_*b*_) with the rest having *h*^2^∑_*g*_/(2*M*_*s*_). Here *M*_*s*_ = *M* − *M*_*b*_ and we consider *M*_*b*_ = 200 in our numerical studies. Hence a small subset of SNPs will have relatively large effect sizes, while the majority of them have modest effect sizes.

We consider *h*^2^ = 0.1, 0.3, 0.5 in the simulations, and compare two approaches for estimating ∑: (1) the sample correlation matrix of summary Z-statistics (excluding genome-wide significant SNPs), denoted as 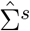; (2) the proposed robust LD score regression, denoted as 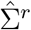; The LD scores are pre-computed based on the 1000 Genomes Project European samples (Abecasis *et al.*, 2012). The LD score regression of Bulik-Sullivan et *al.* (2015*a*) performs slightly worse than the robust LD score regression for estimating both trait and genetic correlations. We provided complete results at the supplementary materials section 3.1. Table 1 summarizes the bias and RMSE computed over 100 simulations. The robust LD score regression based approach performs much better than the naive sample correlation based estimates, which had much larger biases. The biased trait correlation estimates can lead to inflated or conservative type I errors for the down-stream multi-trait association tests (see Table 3 in section 3.1 at the supplementary materials).

**Table 1:**
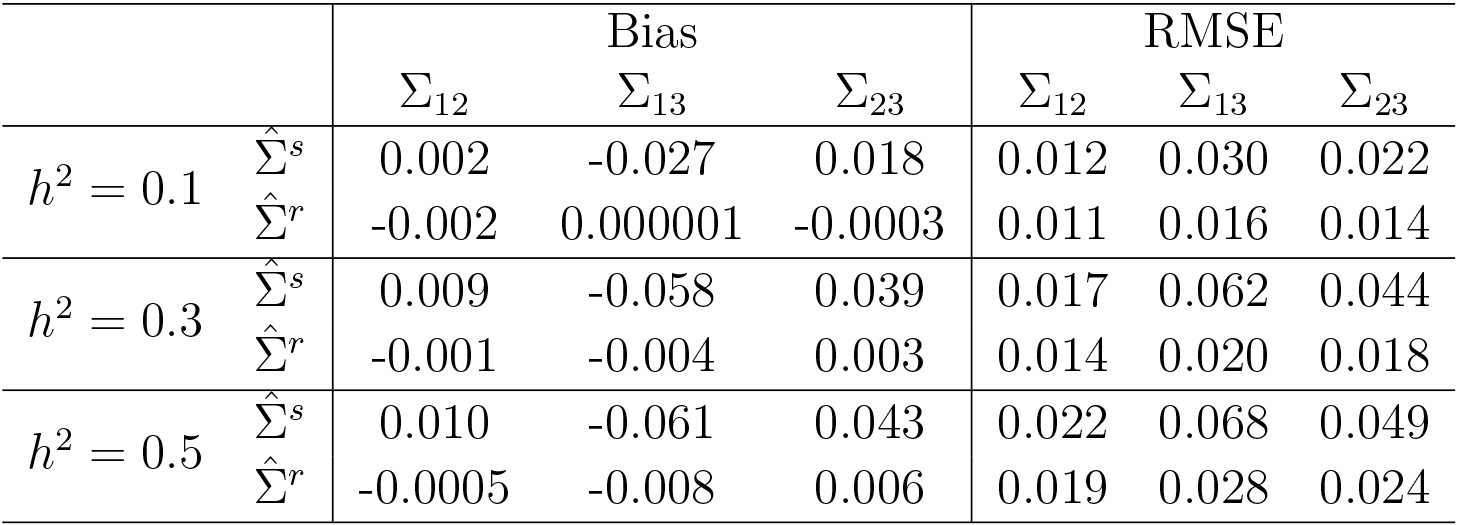
Bias and RMSE of estimating marginal correlations ∑.

### 3.2 Multi-trait association test

We evaluate the performance of proposed tests compared to the following tests: (1) the minimum marginal p-value across traits (denoted as minP); (2) the sum of Z-statistics (denoted as SZ) along the vein of fixed effects meta-analysis; (3) the sum of squared Z-statistics (denoted as SZ2) along the vein of heterogeneity effects meta-analysis. These three tests do not explicitly account for the trait correlations, and generally have less favorable performance. We can compute their analytical p-values very efficiently. The PC based test performs well when the top PC captures majority of the association signals. The SZ test has good performance when all marginal trait effects follow the same direction. In contrast, both *Q* and SZ2 are quadratic tests and have more robust performance. The adaptive test AT has very robust and consistent performance.

We first evaluate the type I errors of proposed tests by simulating 10^10^ random vectors from 𝒩(0, ∑), where ∑ is a correlation matrix with ∑_12_ = −0.11, ∑_13_ = −0.36, ∑_23_ = 0.23. All proposed tests have well controlled type I errors at the significance levels 10^−6,−7,−8^ (see Table 2).

**Table 2:**
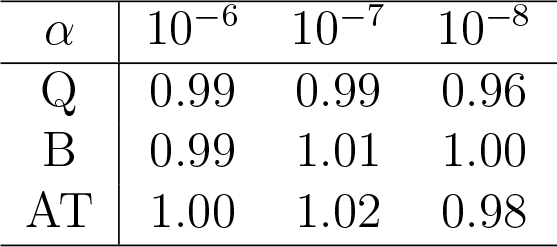
Type I errors divided by the significance level *α* estimated over 10^10^ Monte Carlo simulations: *B* is the PC based test; *Q* is the omnibus chi-square test; AT is the adaptive test.

We evaluate the power of different methods under 5 × 10^−8^ significance level based on simulating 10^7^ random vectors from 𝒩(Δ, ∑). We consider setting Δ as fixed values, and randomly simulating Δ from 𝒩(0, 3) and uniform distributions, *U*(−5, 5), *U*(0, 5). For fixed Δ, we decompose 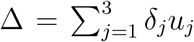 with (*u*_1_, *u*_2_, *u*_3_) being the three eigenvectors of ∑. When the signal vector Δ is completely captured by the first PC (i.e., δ_2_ = δ_3_ = 0), the PC based test *B* has the optimal power. Table 3 shows the estimated power under various settings. The first setting favors *B*, which performs much better than *Q*. However *B* is sensitive to the signal distribution, and performs much worse when the top PC has weak association signal. The minimum p-value based test (minP) performs well when one of the traits dominates the association signal, but otherwise it generally has suboptimal performance. The SZ test performs well when all marginal effects follow the same directions. The SZ2 test is relatively more robust and performs well with multiple large marginal effects. In contrast, both *Q* and AT have very robust and consistent performance over all settings. The adaptive test AT can truly combine the strength of both *Q* and *B* and outperform both tests when they have comparable powers.

**Table 3:** Power (%) under 5 × 10^−8^ significance level: *B* is the PC based test; *Q* is the omnibus chi-square test; AT is the adaptive test; minP is the minimum p-value based test, SZ is the sum of Z-statistics, and SZ2 is the sum of squared Z-statistics. Data simulated from 𝒩(Δ, ∑), where 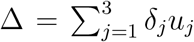 with (*u*_1_, *u*_2_, *u*_3_) being the three eigenvectors of ∑.

| $\Delta$ | $(\delta_1, \delta_2, \delta_3)$ | minP | SZ | SZ2 | B | Q | AT |
| --- | --- | --- | --- | --- | --- | --- | --- |
| (3.59,-2.71,-3.97) | (6,0,0) | 6.45 | 0.02 | 28.65 | 30.06 | 16.93 | 26.42 |
| (2.56,-4.42,-3.73) | (6,2,0) | 13.18 | 2.66 | 38.16 | 30.06 | 29.72 | 34.10 |
| (4.82,-3.25,-2.49) | (6,0,2) | 21.14 | 0.00 | 38.03 | 30.06 | 36.40 | 38.98 |
| (5.33,-2.40,-2.61) | (6,-1,2) | 37.83 | 0.00 | 40.57 | 30.06 | 40.12 | 41.85 |
| (4.63,-0.83,-2.81) | (5,-2,1) | 15.59 | 0.00 | 15.36 | 8.93 | 13.68 | 14.40 |
| (3.22,2.31,-3.21) | (3,-4,-1) | 1.50 | 0.00 | 7.05 | 0.14 | 19.98 | 16.59 |
| (1.52,-4.26,1.26) | (2,3,3) | 8.27 | 0.00 | 2.25 | 0.01 | 25.21 | 21.25 |
| (5,0,0) | (2.99,2.55,-3.09) | 25.99 | 1.07 | 5.31 | 0.14 | 29.78 | 25.48 |
| (2.71,-5.16,-0.07) | (4,3,3) | 31.59 | 0.00 | 21.51 | 1.52 | 51.86 | 46.86 |
| (-0.49,-2.71,-4.63) | (4,2,-3) | 15.56 | 30.10 | 11.76 | 1.52 | 33.28 | 29.07 |
| $\Delta \sim U(0, 5)$ | | 9.15 | 35.23 | 13.59 | 0.17 | 39.29 | 36.66 |
| $\Delta \sim U(-5, 5)$ | | 9.08 | 10.11 | 13.92 | 2.92 | 33.23 | 31.53 |
| $\Delta \sim \mathcal{N}(0, 3)$ | | 20.67 | 11.13 | 20.12 | 4.04 | 34.28 | 32.85 |

We have conducted more simulation studies investigating the performance of different methods under various settings of different number of traits and trait dependence. The complete results are available at the supplementary materials section 4.

## 4 Application

We conduct comprehensive analysis of GWAS meta-analysis results for multiple lipids and glycemic traits. The proposed tests have performed better than the other three competing methods (minP, SZ and SZ2). In the following, we mainly focus on results for the proposed methods, and leave the complete results to the supplementary materials. We note that it is more productive to treat the proposed tests as a complementary approach to the existing single-trait based test. We thus present joint association test results excluding genome-wide significant SNPs for any trait. The analysis results including all SNPs are provided at the supplementary materials section 5.

### 4.1 Analysis of lipids GWAS results

We analyze the GWAS meta-analysis results for three plasma lipids (LDL cholesterol, triglyceride and total cholesterol) based on around 100,000 European individuals from the Global Lipids Consortium (Teslovich *et al.*, 2010). We note that the Global Lipids Consortium conducted a followup study using a Metabochip with a small panel of preselected SNPs based on 190,000 European samples at Willer *et al.* (2013), which will be used for partial validation in our analysis.

At the 5 × 10^−8^ genome-wide significance level, the omnibus chi-square test *Q* identified 44 significant loci, and the PC based test *B* identified 22 significant loci. The adaptive test AT identified 40 significant loci, including the majority of significant loci identified by *Q* and *B*. Figure 1 (a) and (b) compare the number of significant loci and SNPs identified by the proposed joint tests.

**Figure 1:**
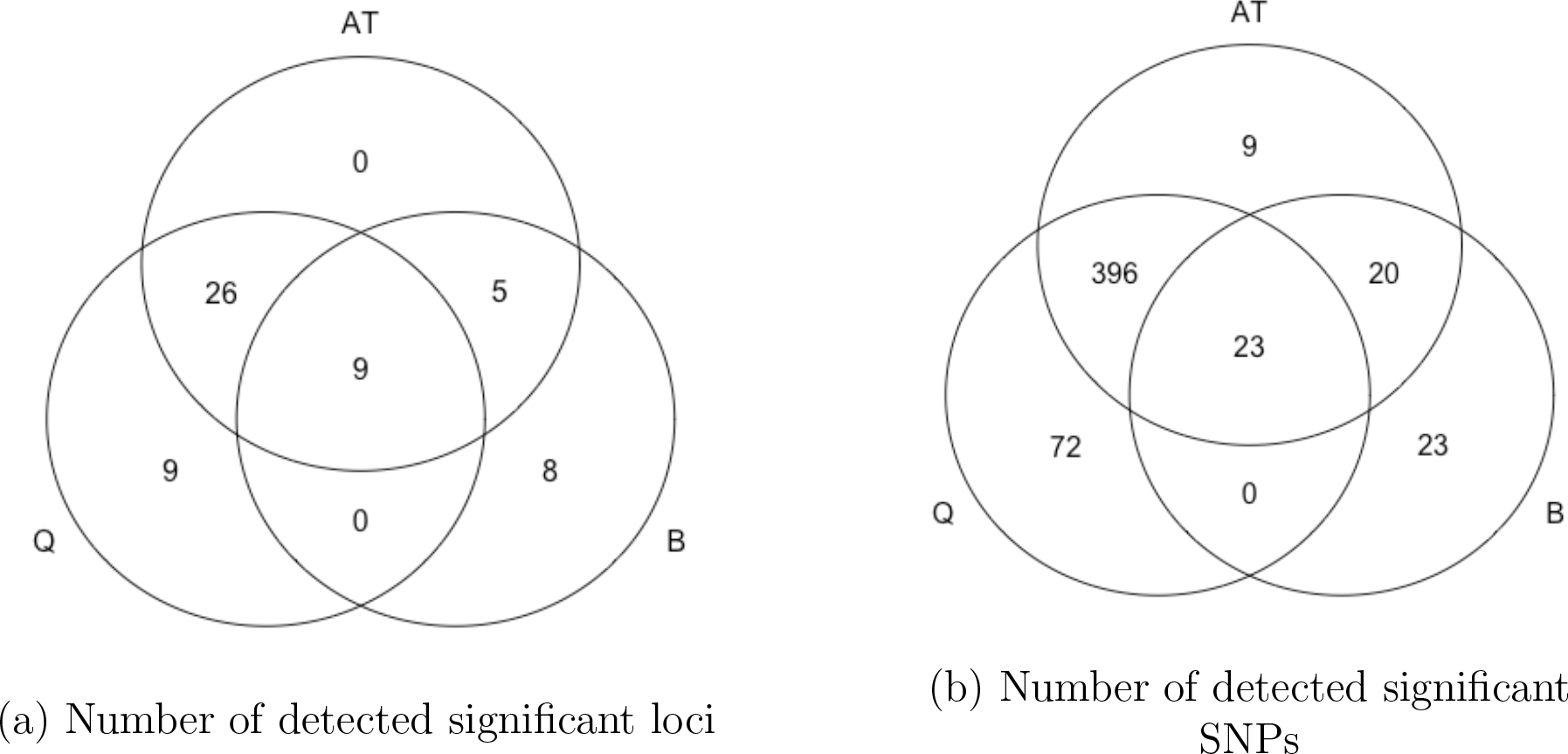
Venn diagram of number of significantly associated loci and SNPs identified by three joint association test methods based on the lipids GWAS summary data.

Many of those identified significant SNPs have been found genome-wide significant in the followup study of Willer *et al.* (2013). Table 4 listed the total number of SNPs and loci identified by three joint association test methods. Table 5 listed the test results for those identified novel loci.

**Table 4:** Number of significant SNPs and loci identified for the lipids GWAS summary data: listed within parentheses are the number of SNPs and loci that have been found genome-wide significant in the followup study of Willer *et al.* (2013).

|  | Q | B | AT | Total |
| --- | --- | --- | --- | --- |
| SNPs | 491 (364) | 66 (38) | 448 (336) | 543 (390) |
| Loci | 44 (37) | 22 (14) | 40 (36) | 57 (43) |

**Table 5:** Novel significant loci identified by the proposed joint test methods: listed are the joint test p-values and the minimum marginal test p-values across all traits in Teslovich *et al.* (2010) and Willer *et al.* (2013) (denoted as minP-2010 and minP-2013 respectively).

| SNP | Chr | Gene | Q | B | AT | minP-2010 | minP-2013 |
| --- | --- | --- | --- | --- | --- | --- | --- |
| rs6730449 | 2 | DYNC2LI1 | 5.75e-07 | 1.72e-08 | 3.72e-08 | 1.83e-07 | 2.43e-06 |
| rs762861 | 4 | HGFAC | 1.52e-07 | 2.74e-08 | 4.72e-08 | 9.15e-07 | 5.04e-07 |
| rs7705104 | 5 | PELO | 2.38e-08 | 9.07e-01 | 5.12e-08 | 1.72e-02 | - |
| rs10462958 | 5 | TIMD4 | 7.02e-07 | 2.99e-08 | 6.42e-08 | 2.03e-07 | - |
| rs499921 | 6 | FRK | 4.99e-07 | 2.70e-08 | 5.80e-08 | 1.81e-07 | 5.07e-08 |
| rs12667186 | 7 | SP4 | 9.69e-07 | 3.14e-08 | 6.75e-08 | 1.36e-07 | 1.08e-07 |
| rs17148062 | 7 | POT1 | 4.66e-08 | 7.65e-01 | 9.98e-08 | 6.14e-02 | - |
| rs4455806 | 8 | SOX17 | 1.29e-06 | 4.60e-08 | 9.85e-08 | 1.84e-07 | 5.09e-08 |
| rs10090964 | 8 | SDCBP | 5.78e-07 | 2.98e-08 | 6.40e-08 | 5.09e-08 | 7.37e-08 |
| rs4573621 | 10 | GPAM | 4.54e-08 | 4.14e-02 | 9.72e-08 | 2.70e-04 | 2.09e-04 |
| rs4238103 | 12 | LARP4 | 4.08e-09 | 9.05e-02 | 4.08e-09 | 3.83e-04 | - |
| rs9600211 | 13 | KLF12 | 1.57e-09 | 1.88e-01 | 1.57e-09 | 1.52e-05 | - |
| rs4419034 | 15 | FRMD5 | 4.87e-08 | 4.30e-01 | 1.04e-07 | 5.05e-08 | 1.51e-06 |
| rs16959082 | 17 | RCVRN | 3.67e-08 | 5.32e-02 | 7.87e-08 | 7.44e-05 | 4.32e-01 |

Table 6 listed several significant SNPs only identified by AT and their minimum p-values across three traits in the Teslovich et *al.* (2010) and Willer *et al.* (2013) study (denoted as minP-2010 and minP-2013 respectively).

**Table 6:** Significant SNPs only identified by AT, but missed by *Q* and *B*: listed are the joint test p-values and the minimum marginal test p-values across all traits in Teslovich *et al.* (2010) and Willer *et al.* (2013) (denoted as minP-2010 and minP-2013 respectively).

| SNP | Chr | Gene | Q | B | AT | minP-2010 | minP-2013 |
| --- | --- | --- | --- | --- | --- | --- | --- |
| rs4808957 | 19 | GATAD2A | 5.05e-08 | 2.05e-07 | 4.52e-08 | 9.58e-07 | 1.83e-14 |
| rs3095340 | 6 | KIAA1949 | 5.37e-08 | 1.75e-07 | 4.61e-08 | 1.27e-07 | 4.23e-07 |
| rs505870 | 6 | SLC22A3 | 7.41e-08 | 5.27e-08 | 4.12e-08 | 3.34e-07 | 1.05e-09 |
| rs10806731 | 6 | LPA | 6.16e-08 | 1.13e-07 | 4.75e-08 | 6.40e-07 | - |

### 4.2 Analysis of GWAS results for glycemic traits

We also analyze the GWAS meta-analysis results for two glycemic traits: fasting glucose (FG) and indices of *β*-cell function (HOMA-B) based on 46,186 non-diabetic European samples conducted by the international MAGIC consortium (Dupuis *et al.*, 2010). Figure 2 (a) and (b) show the venn diagram for the total number of identified significant loci and SNPs by the proposed joint association test methods at the 5 × 10^−8^ genome-wide significance level. The chi-square test *Q* identified 8 significantly associated loci, and the PC test *B* identified 13 significant loci. The adaptive association test AT identified all significant loci identified by *Q* and 4 additional significant locus identified by *B*.

**Figure 2:**
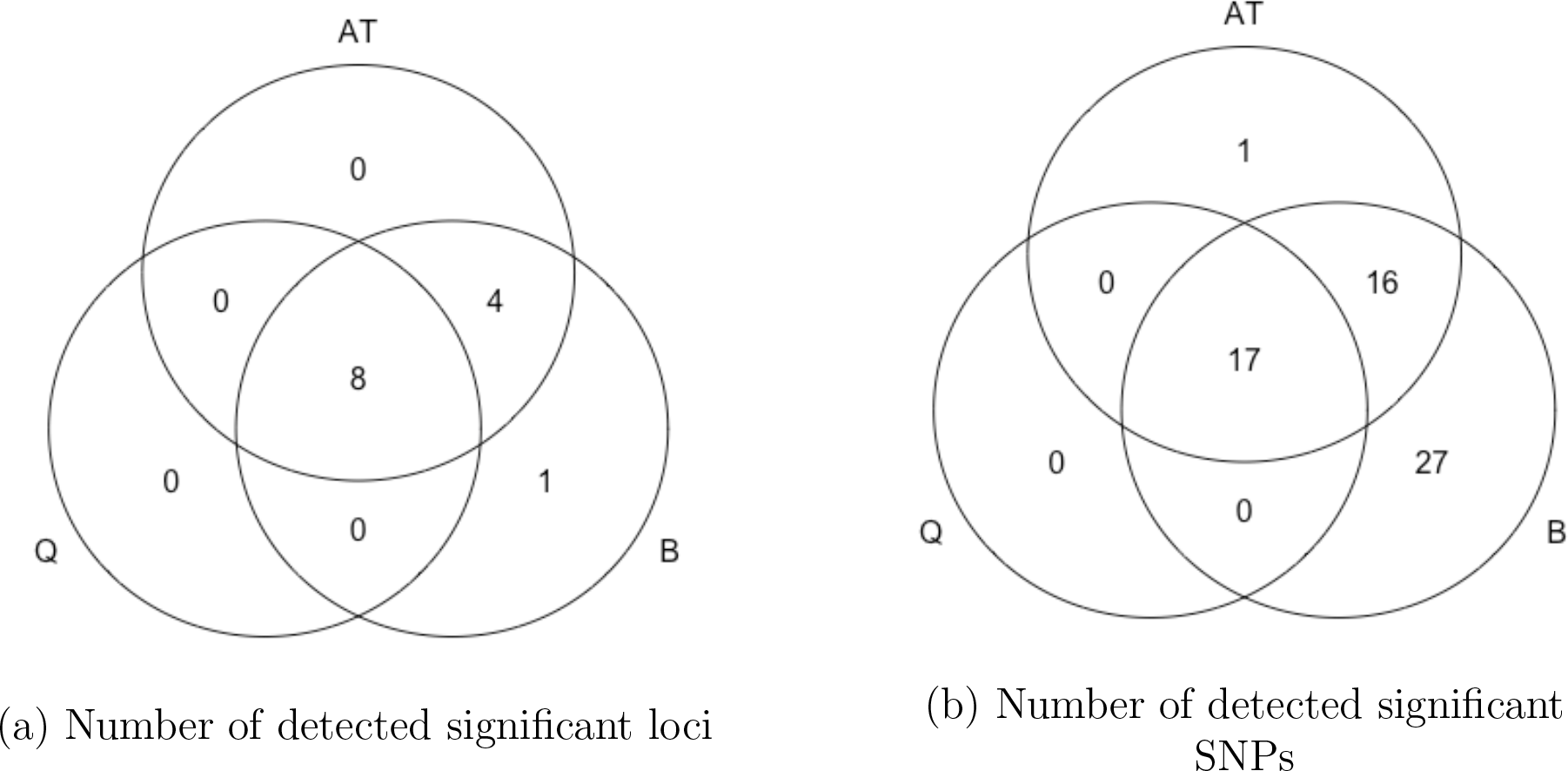
Venn diagram of the total number of significant loci and SNPs identified by three multi-trait test methods based on glycemic GWAS summary data

The comprehensive analysis results of the GWAS summary data for the lipids and glycemic traits are provided at the supplementary materials section 5.

## 5 Discussion

Many GWAS have been conducted in the past decade and successfully identified thousands of diseases related variants. However there are still significant missing heritability for most traits and there remain many more genetic variants with small effects to be discovered. In the post GWAS era, owing to many difficulties of sharing raw genotype and phenotype data, it is useful to develop statistical methods that can leverage the publicly available GWAS summary data to identify more novel genetic variants. In this paper, we have focused on testing SNP association across multiple traits, which has been shown to have improved test power than individual trait based association test. To properly control the false positives for multi-trait association test, we need to accurately estimate the across trait correlations. Our results show that the commonly used approach of using empirical correlation matrix of summary Z-scores should be avoided if possible, since it can produce highly biased estimates and lead to significantly inflated type I errors for polygenic traits. The proposed LD score regression based approach could produce more accurate estimates, and has performed well in our numerical studies. In GWAS we typically need to test tens of millions of SNPs, and it is desirable to develop efficient statistical methods. All our proposed methods are scalable to genome-wide association test: we can quickly compute their analytical p-values without the need of resampling or permutation.

In this paper, we have mainly focused on those efficient and genome-wide scalable joint association test methods with analytically computed p-values and proper control of type I errors, and haven’t studied those methods that often require computationally intensive Monte Carlo or MCMC simulations (see, e.g., Stephens, 2013; Kim *et al.*, 2015; Shim et al., 2015).

Besides combining multiple traits to boost overall association test power, it also helps to further integrate multiple variants in a gene or pathway. Individually these variants may have weak effects, but in combination they could exert large effects on the outcome. It is worthwhile to develop efficient and powerful “multi-trait multi-variant association test” methods using just the GWAS summary data (Cichonska et al., 2016; Zhu and Stephens, 2017).

We have implemented the proposed methods in an R package publicly available online. The supplementary materials contain the detailed analysis results for the lipids and glycemic GWAS summary data, and sample codes to install and use the developed R package.

## Acknowledgements

We are grateful to the University of Minnesota Supercomputing Institute for assistance with the computations. This work was supported in part by NIH grant GM083345 and CA134848.

